# Spatiotemporal Characterization of Cyclooxygenase Pathway Enzymes During Vertebrate Embryonic Development

**DOI:** 10.1101/2024.08.02.606390

**Authors:** Tess A. Leathers, Raneesh Ramarapu, Crystal D. Rogers

## Abstract

Vertebrate development is regulated by several complex well-characterized morphogen signaling pathways, transcription factors, and structural proteins, but less is known about the enzymatic pathways that regulate early development. We have identified that factors from the inflammation-mediating cyclooxygenase (COX) signaling pathway are expressed at early stages of development in avian embryos. Using *Gallus gallus* (chicken) as a research model, we characterized the spatiotemporal expression of a subset of genes and proteins in the COX pathway during early neural development stages. Specifically, here we show expression patterns of COX1, COX2, and microsomal prostaglandin E synthase-2 (mPGES-2) as well as the genes encoding these enzymes. Unique expression patterns of individual players within the COX pathway suggest that they may play non-canonical/non-traditional roles in the embryo compared to their roles in the adult. Future work should examine the function of the COX pathway in tissue specification and morphogenesis and determine if these expression patterns are conserved across species.

Graphical abstract.Spatiotemporal characterization of cyclooxygenase pathway enzymes during vertebrate embryonic development.

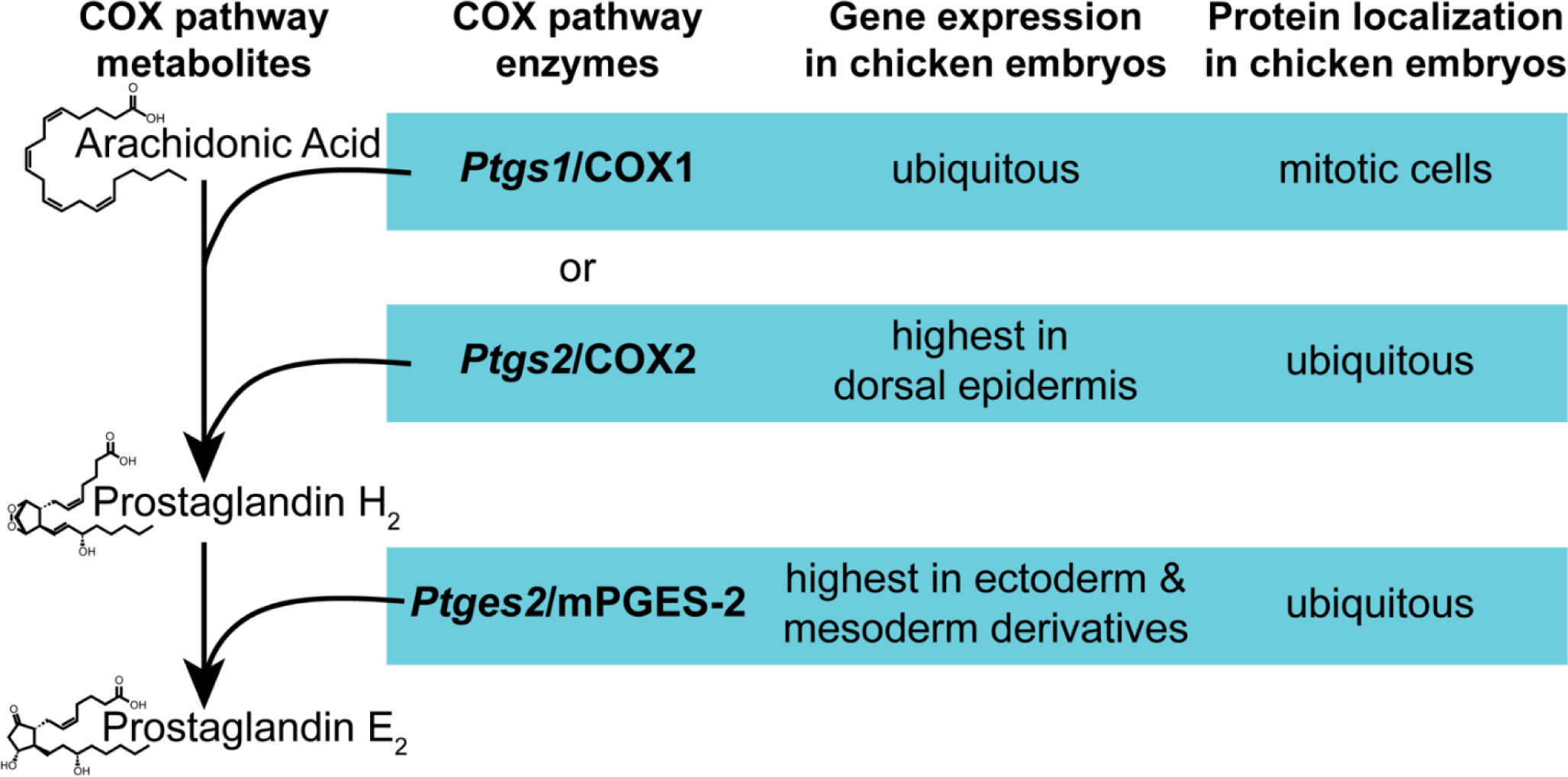

## Introduction

Perturbation of the cyclooxygenase (COX) pathway during pregnancy is linked to developmental anomalies, but little is known about its typical localization and function in the embryo (1, 2). Inhibition of COX activity during embryogenesis using nonsteroidal anti-inflammatory drugs increases the risk of neural tube defects (3), abnormal cardiogenesis (4), craniofacial clefts (3), and impaired gut innervation (5) among other issues (1, 2). Meanwhile, after infection or injury, the COX pathway can be upregulated by multiple cytokines (6–9), which are linked to a predisposition for several neuropathologies when activated *in utero* (10). COX pathway signaling has been implicated in uterine implantation (11), angiogenesis (9, 12, 13), formation of the central and enteric nervous systems (3, 5), skeletal development (2), and immune system modulation (14). Because of its wide-ranging implications, defining the tissues in which COX pathway factors are expressed in the embryo is a crucial first step in understanding how changes in the pathway will affect development.

Isoenzymes COX1 and COX2 catalyze the first step in the biosynthesis of a variety of inflammation-mediating signaling molecules called prostaglandins (PG) and thromboxanes (TX), collectively known as prostanoids. Specifically, after arachidonic acid is freed from phospholipids by phospholipase A_2_ enzymes (PLA_2_), COX isoenzymes convert the long chain fatty acid into PGG_2_, then reduce PGG_2_ into PGH_2_ (15). It is because of this activity that COX isoenzymes are also called prostaglandin-endoperoxide synthases and the official gene names of COX1 and COX2 are *Ptgs1* and *Ptgs2*, respectively. COX1 is traditionally referred to as the “housekeeping” COX isoform and is broadly expressed in adult human tissues (16). Subcellularly, it is associated with the endoplasmic reticulum and nuclear envelope (17, 18). However, studies during embryonic development suggest that *Ptgs1* abundance may be more spatiotemporally dynamic than in the adult (19–21). In contrast, COX2 is traditionally seen as an inducible isoenzyme, responding to illness or injury (17, 22). However, some studies have found COX2 to be widely distributed in adult human tissues (16). Like COX1, COX2 is localized to the endoplasmic reticulum and nuclear envelope, but while COX1 is equally distributed, in murine 3T3 cells and human and bovine endothelial cells, COX2 was twice as concentrated in the nuclear envelope than the endoplasmic reticulum (18). *Ptgs2* was found at elevated levels in fetal rat tissue from gestation days 17-20 but was not detected in embryonic tissues at gestational days 7-13 (23). In zebrafish embryos, *Ptgs2* was seen as early as two-somite stage in the anterior neuroectoderm (20).

Downstream of COX1 and COX2, PGH_2_ is further metabolized by terminal prostanoid synthases like microsomal PGE synthase-2 (mPGES-2). mPGES-2 is one of three PGE synthases responsible for producing PGE_2_ (24). There are two membrane-associated forms, mPGES-1 and mPGES-2, and a cytosolic form, cPGES (24). mPGES-2 is encoded by the *Ptges2* gene and functions independently of glutathione, unlike the other PGES enzymes (24, 25). In HEK293 and BEAS-2B cells, mPGES-2 is first synthesized as a Golgi membrane-associated protein then found in the cytoplasm once its N-terminal hydrophobic domain is removed (27). At the tissue level, mPGES-2 is reported in the brain, heart, skeletal muscle, kidney, and liver of adults (24). The gene encoding mPGES-1, *Ptges*, is reported to be expressed as early as gastrulation stages (20) and blocking its translation prevents normal gastrulation movements in zebrafish (26). However, *Ptges2* expression patterns in the embryo are unknown.

Currently, we lack the spatiotemporal expression data for COX pathway enzymes needed to understand the mechanistic role of this pathway at key embryonic stages. Here, using *in situ* hybridization chain reaction (HCR), immunohistochemistry (IHC), and molecular staining, we visualize the expression of three COX pathway enzymes during neurulation in *Gallus gallus* (chicken) embryos (28). We visualize the expression of *Ptgs1*, *Ptgs2*, *Ptges2* transcripts, and the proteins encoded by each gene (Table 1) to characterize their tissue-specific localization during neural tube closure and fusion. Our results demonstrate that COX pathway enzymes are dynamically and broadly expressed in neurulating amniotic embryos. Additionally, while COX2 appears to be ubiquitously expressed in all cell types, COX1 and mPGES have unique tissue and subcellular specific localization.

**Table 1.** COX pathway enzymes characterized in this study.

| Protein Name | Gene Name | NCBI Reference Sequence | Aliases |
| --- | --- | --- | --- |
| Cyclooxygenase 1 (COX1) | *Ptgs1* | XM_040685541.2 | Prostaglandin-Endoperoxide Synthase 1, Prostaglandin G/H Synthase 1, Prostaglandin H<sub>2</sub> Synthase 1 |
| Cyclooxygenase 2 (COX2) | *Ptgs2* | NM_001167719.2 | Prostaglandin-Endoperoxide Synthase 2, Prostaglandin G/H Synthase 2, Prostaglandin H<sub>2</sub> Synthase 2 |
| Microsomal Prostaglandin E Synthase-2 (mPGES-2) | *Ptges2* | XM_040685892.2 | Membrane-Associated Prostaglandin E Synthase-2 |

## Results

### COX pathway gene expression across early developmental stages

To begin our analysis of COX pathway factor gene expression, we combined two open-source single cell RNA-sequencing datasets spanning chicken stages Hamburger Hamilton (HH) stage 4 - HH9. Clusters were defined through unsupervised clustering and were annotated using published marker genes and gene sets (Figure 1A,1B, Supplementary Figure 1A-1M). Following clustering, we investigated COX pathway members across the datasets and identified that a majority are expressed at low levels across many tissues. (Figure 1, Supplementary Figure 2). Specifically, *Ptgs1*, which encodes COX1, was more highly expressed in the transitional neural plate and neural tube (TNP/NT) and the non-neural ectoderm (NNE) regions (Figure 1C, 1D). Over developmental time, the TNP/NT will become the brain and spinal cord while the NNE will become sensory placodes and epidermal tissues. The *Ptgs2* gene, which encodes COX2 protein, was expressed at significantly lower levels across ectodermally-derived tissues but had increased expression in the anterior lateral plate mesoderm (ALPM) cells (Figure 1E, 1F). In contrast to either *Ptgs1* or *Ptgs2*, *Ptges2*, which encodes mPGES-2, has much broader expression than the upstream pathway enzymes. The definitive neural (Def Neu) populations had the highest *Ptges2* expression, but the gene was expressed at detectable levels in all cell types with the exception of the posterior neural plate/neural tube (PNP/NT) (Figure 1G, 1H).

**Figure 1.**
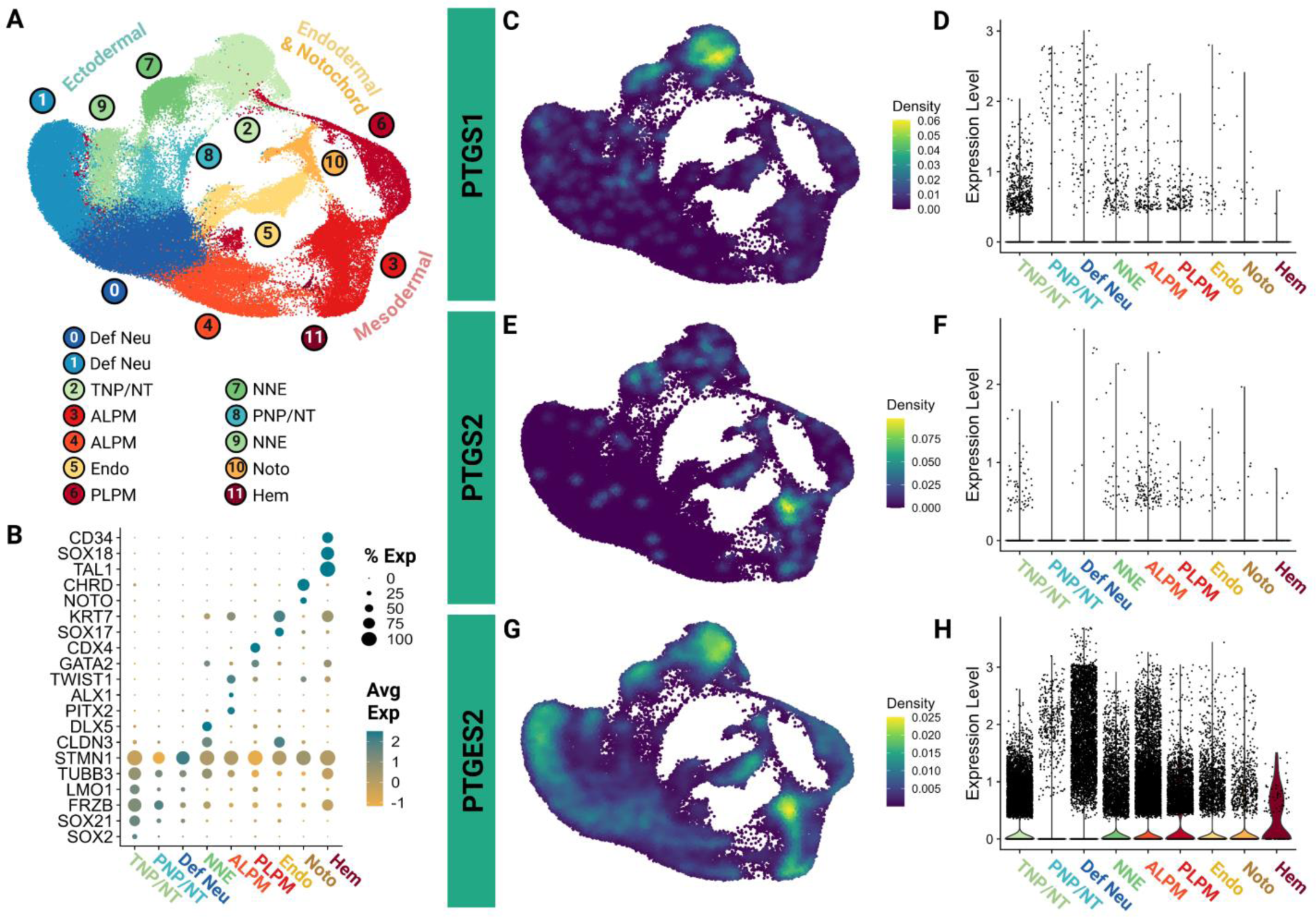
Analysis of publicly available scRNA-sequencing data of chick embryos shows cell type specific expression of select cyclooxygenase pathway isoenzymes. (A) UMAP demonstrating the unsupervised clustering results of scRNA-sequencing from whole chick embryos. (B) Dot plot demonstrating cell type specific marker expression of the 9 major cell types identified across the 12 clusters. (C,E,G) Feature UMAPs showing gene expression of select enzymes by kernel density estimation. (D,F,H) Violin plots demonstrating expression of select enzymes across the 9 major cell types identified. Definitive Neural lineage, Def Neu; Transitional neural plate/ neural tube, TNP/NT; Anterior lateral plate mesoderm, ALPM; Endoderm, Endo; Posterior lateral plate mesoderm, PLPM; Non-neural ectoderm, NNE; Posterior neural plate/ neural tube, PNP/NT; Notochord, Noto; Hematogenic cells, Hem.

### COX1 transcript and protein expression during neurulation

Previous studies suggested that *Ptgs1* is ubiquitously expressed during embryonic stages (20, 21) and becomes more spatially restricted as development progresses (19), but *Ptgs1* gene and subsequent COX1 protein spatial expression has not yet been characterized at these early developmental stages in amniotic embryos over the course of their development. To spatially visualize the *Ptgs1* gene expression identified in single-cell analysis (Figure 1) during neurulation, we used HCR in chicken embryos at stages HH7-10 in conjunction with a DAPI stain to visualize nuclei. We provide a color-coded schematic map of HH9 chicken embryo sections to delineate specific tissues that express each factor (Figure 2). We paired *Ptgs1* probes with a probe for neural cadherin (*Ncad*) as a marker of neural tube and paraxial mesoderm because single cell analysis (Figure 1) suggested that *Ptgs1* would be expressed robustly in the neural tube compared to other tissues. We observed that *Ptgs1* transcript appears to be expressed ubiquitously throughout the embryo from HH7-10 and its expression does overlap with *Ncad* in the neural tube and paraxial mesoderm (Figure 3A-3H).

**Figure 2.**
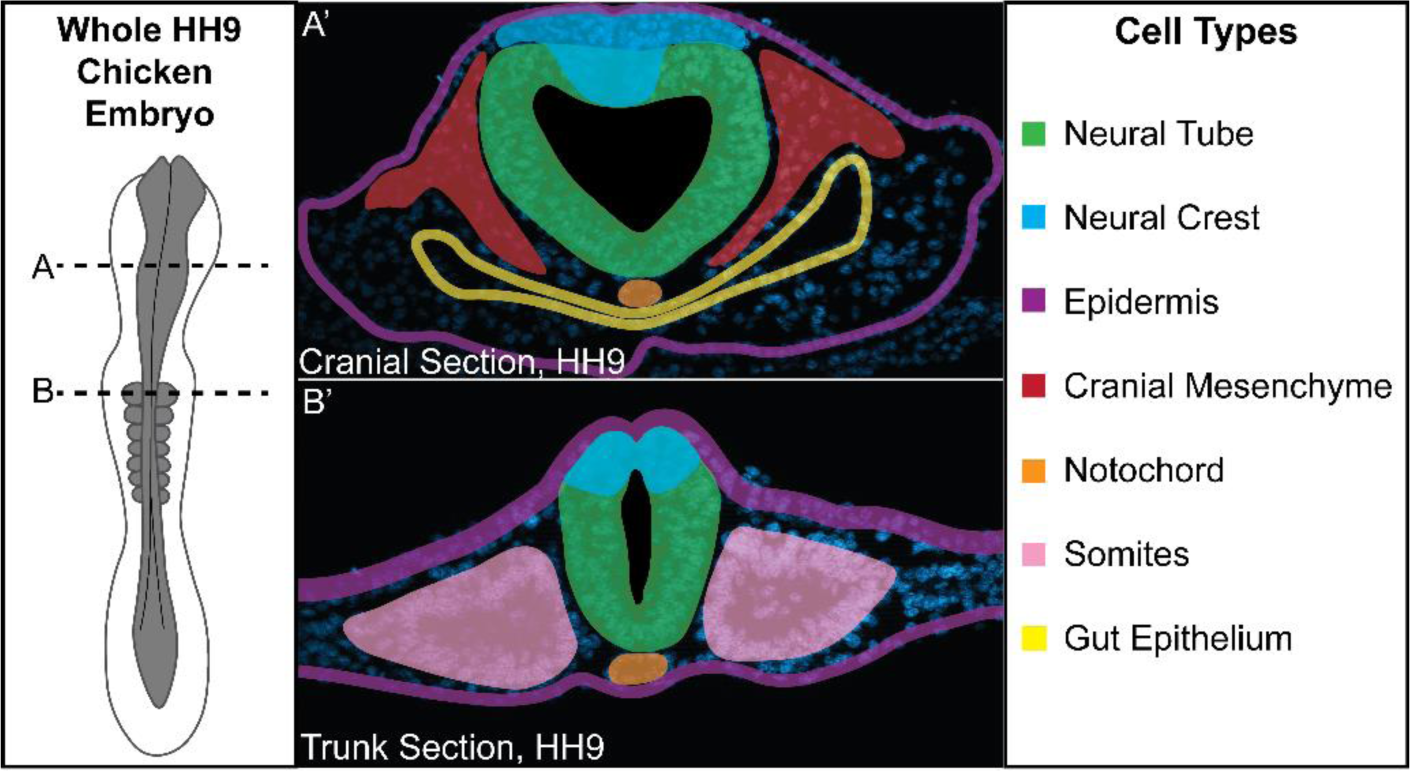
Schematic of cell and tissue types from all three germ layers in a chicken embryo. Whole HH9 chicken embryo illustration shown with transverse cryosections taken at indicated axial levels (A, B). (A’) A cranial transverse section of an HH9 chicken embryo at the axial position indicated by (A) shows cells from the ectodermal (neural crest, neural tube, epidermis), mesodermal (cranial mesenchyme, notochord), and endodermal (gut epithelium) lineages. (B’) A trunk transverse section of an HH9 chicken embryo at the axial position indicated by (B) shows the same cell types in the trunk with the addition of somites from the mesodermal lineage and lack of gut epithelium and cranial mesenchyme due to the more posterior axial level.

**Figure 3.**
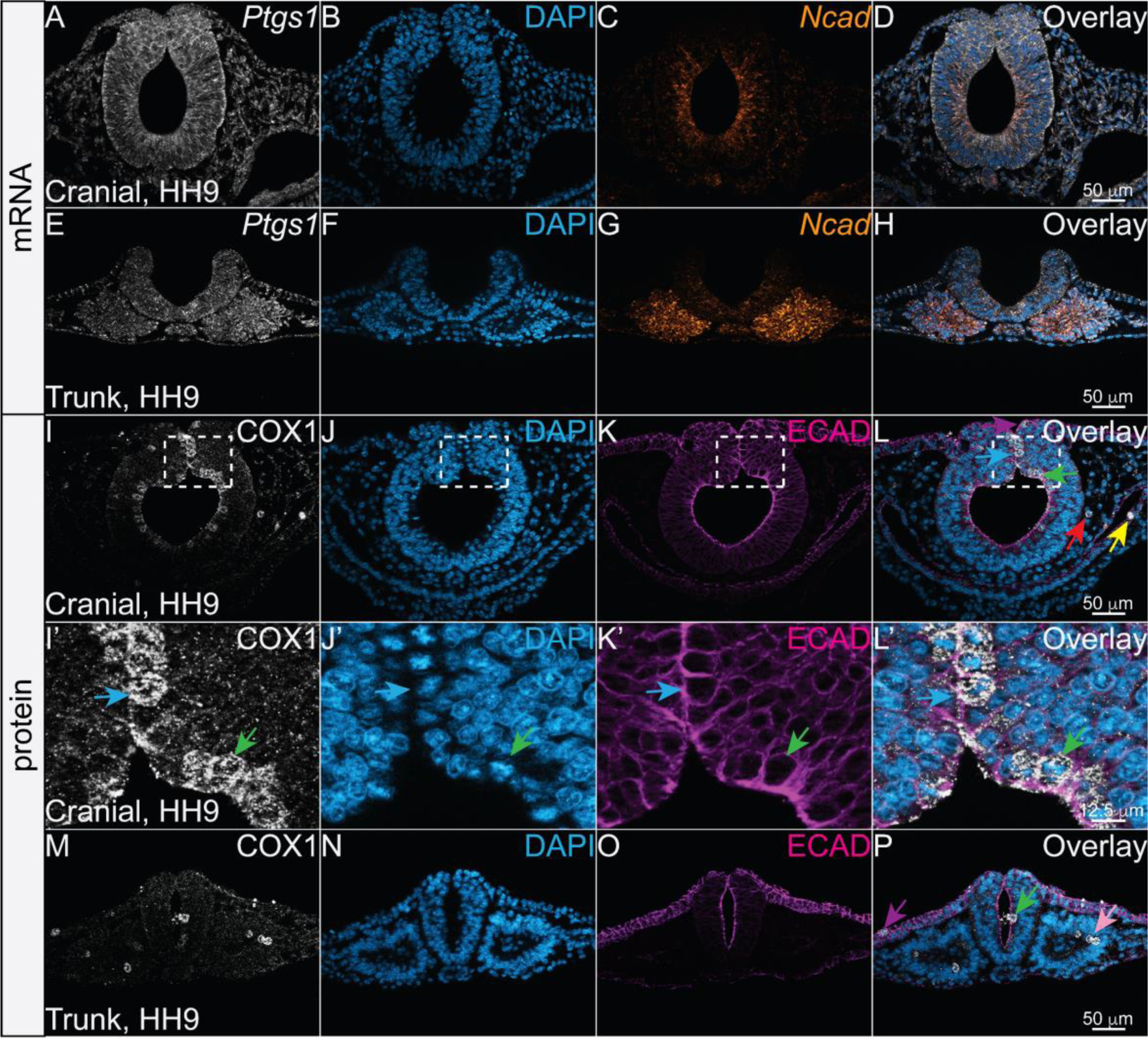
COX1 is localized to mitotic cells in all three germ layers while its transcript is more broadly expressed. (A-H) Cranial and trunk transverse cryosections of an HH9 chicken embryo with HCR showing COX1 transcript *Ptgs1* (white), the nuclear stain DAPI (blue), neural cadherin transcript *Ncad* (orange), and the overlay of all three channels (D, H). *Ptgs1* appears expressed throughout the embryo. (I-P) Cranial and trunk transverse cryosections of an HH9 chicken embryo after IHC using antibodies against the isoenzyme COX1 (white), the nuclear stain DAPI (blue), and epithelial cadherin, ECAD (magenta). (I’-L’) Zoom in on region outlined in (I-L). (I-P) Arrows indicate COX1+ mitotic cells with colors coordinating to the cell or tissue types described in Figure 2. COX1+ mitotic cells are found in ectodermal derivatives (neural tube in green, neural crest in blue, and epidermis in purple), mesodermal derivatives (cranial mesenchyme in red and somites in pink), and endodermal derivatives (gut epithelium in yellow). Scale bar for each row in the last image of the row.

To characterize COX1 protein expression and localization, we performed IHC using an antibody against COX1 paired with epithelial cadherin (ECAD), which localizes to epithelial cell membranes and is broadly expressed, and DAPI stain in chicken embryos at stages HH7-10. In contrast to the ubiquitous *Ptgs1* transcripts, the COX1 protein appeared to be specifically upregulated in cells undergoing mitosis across all axial levels and in cells derived from all three germ layers (Figure 3I-3P). In cells derived from the ectoderm, COX1 was seen in mitotic cells of the neural tube, neural crest, and epidermis (Figure 3I-3P, green, blue, and purple arrows).

During early neural development and prior to cortical histogenesis, nuclei from neuroepithelial cells undergo interkinetic nuclear migration and migrate to the apical side of the neural tube to proliferate (29). COX1-positive cells are specifically localized to the apical side of the neural tube and DAPI staining confirms that these cells are in different stages of mitosis (Figure 3I’-3L’, blue and green arrows). Similarly, in the mesodermally-derived cells, COX1 was detected in cranial mesenchyme cells undergoing mitosis (Figure 3I-3L, red arrow). In the endodermally derived cells, COX1 was detected in mitotic cells lining the developing gut (Figure 3I-3L, yellow arrow). Mesodermally-derived somites also undergo interkinetic nuclear migration while proliferating (30), and in trunk sections, COX1 is upregulated in the apical side of the developing somites in cells undergoing mitosis (Figure 3M-3P, pink arrow). Among the various cell types, COX1 is expressed in the cytosol of mitotic cells, and is mutually exclusive from DAPI-stained DNA throughout the stages of mitosis (Figure 3I-3P).

To characterize the expression and subcellular localization of COX1 during each phase of mitosis, we paired COX1 with Alpha Tubulin to mark microtubules and DAPI to mark the chromosomes. COX1 first appears in cells during prophase and is expressed at lower levels by telophase (compare prophase and metaphase to anaphase and telophase, Figure 4A-4P). COX1 appears mutually exclusive from the DAPI signal and may localize to the cytoplasm in a perinuclear location (Figure 4A-4D, 4E-4H). During anaphase, COX1 protein signal weakens and by the end of telophase, COX1 signal appears absent or at low levels in non-dividing cells (Figure 4A-4P, 3I-3P). Notably, COX1 is more distinctly visualized with 2% TCA fixation compared to 4% PFA fixation (Supp. Figure 3) as has been identified for other cytoplasmic and structural proteins (33).

**Figure 4.**
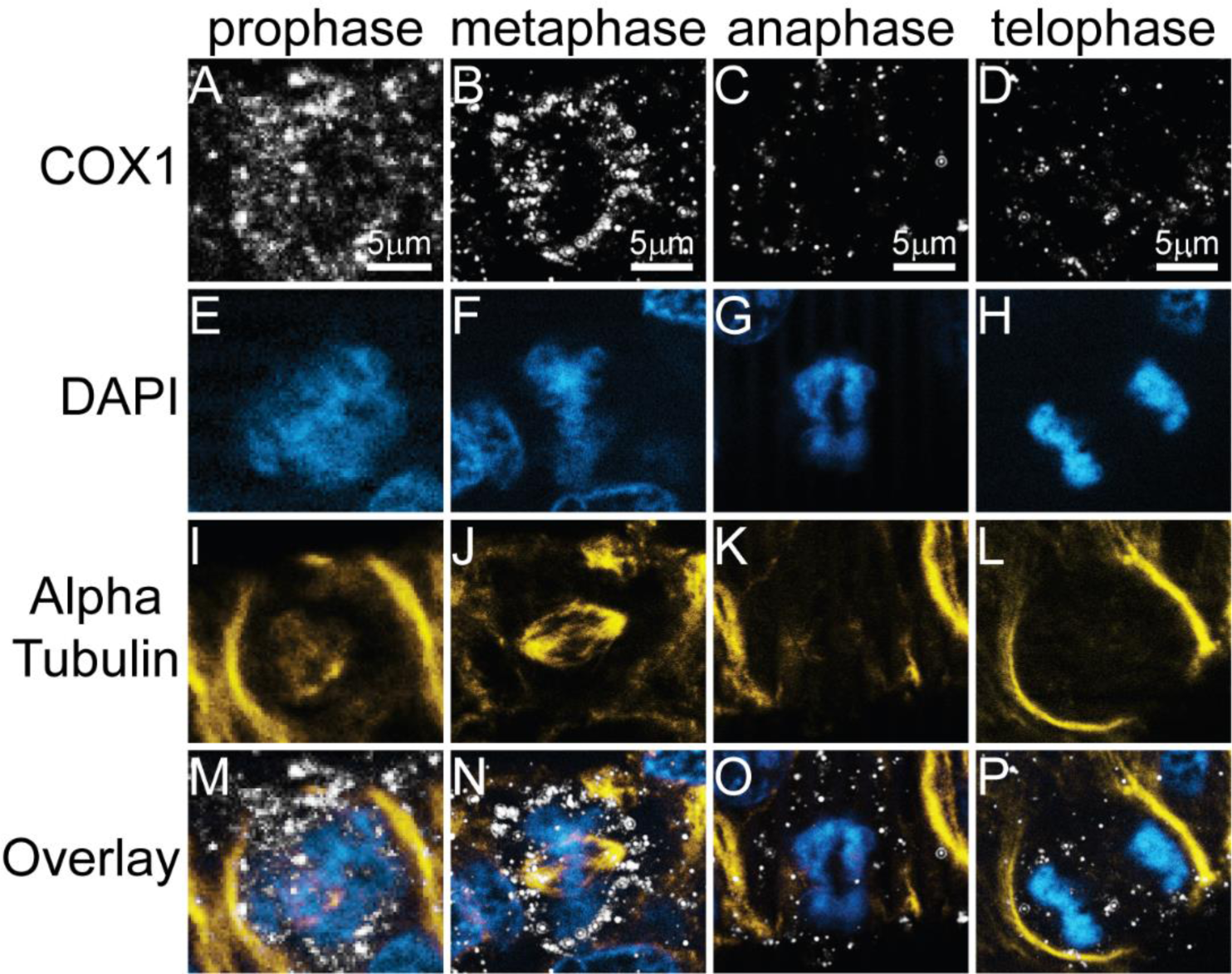
COX1 protein is upregulated in cells undergoing mitosis in the chicken embryo. IHC using antibodies against (A-D) the isoenzyme COX1 (white), (E-H) the nuclear stain DAPI (blue), and (I-L) the microtubule alpha tubulin subunit (yellow) in HH9 and HH10 chicken embryos shows that COX1 is present in cells at each stage of mitosis. Overlays of all three channels shown in (M-P). Scale bar for each column in the first image of the column.

### COX2 transcript and protein expression during neurulation

Based on analysis of scRNA-sequencing datasets, *Ptgs2* expression is low, but existent, in multiple embryonic tissues. COX2 mRNA and protein expression in rat fetal tissues were reported to start at 15 days of gestation (23), but in zebrafish *Ptgs2* was seen as early as the two-somite stage, when organogenesis is just beginning (20, 31). To visualize *Ptgs2* transcript expression during neurulation, we used HCR in chicken embryos at stages HH7-10. We observed that from HH7-8, it is difficult to visualize *Ptgs2* expression due to its low transcript levels (Figure 5A-5C’). At HH9, *Ptgs2* transcript becomes more distinctly expressed in the forebrain and the trunk epidermis with reduced expression in the midbrain (Figure 5D-5F’, purple arrows). In the trunk cryosections, Ptgs2 expression is apparent in multiple tissues, but it appears most highly expressed in dorsal tissues, particularly at the joining of the neural folds, and in the superficial epidermis (Figure 5D-5F’, purple arrows).

**Figure 5.**
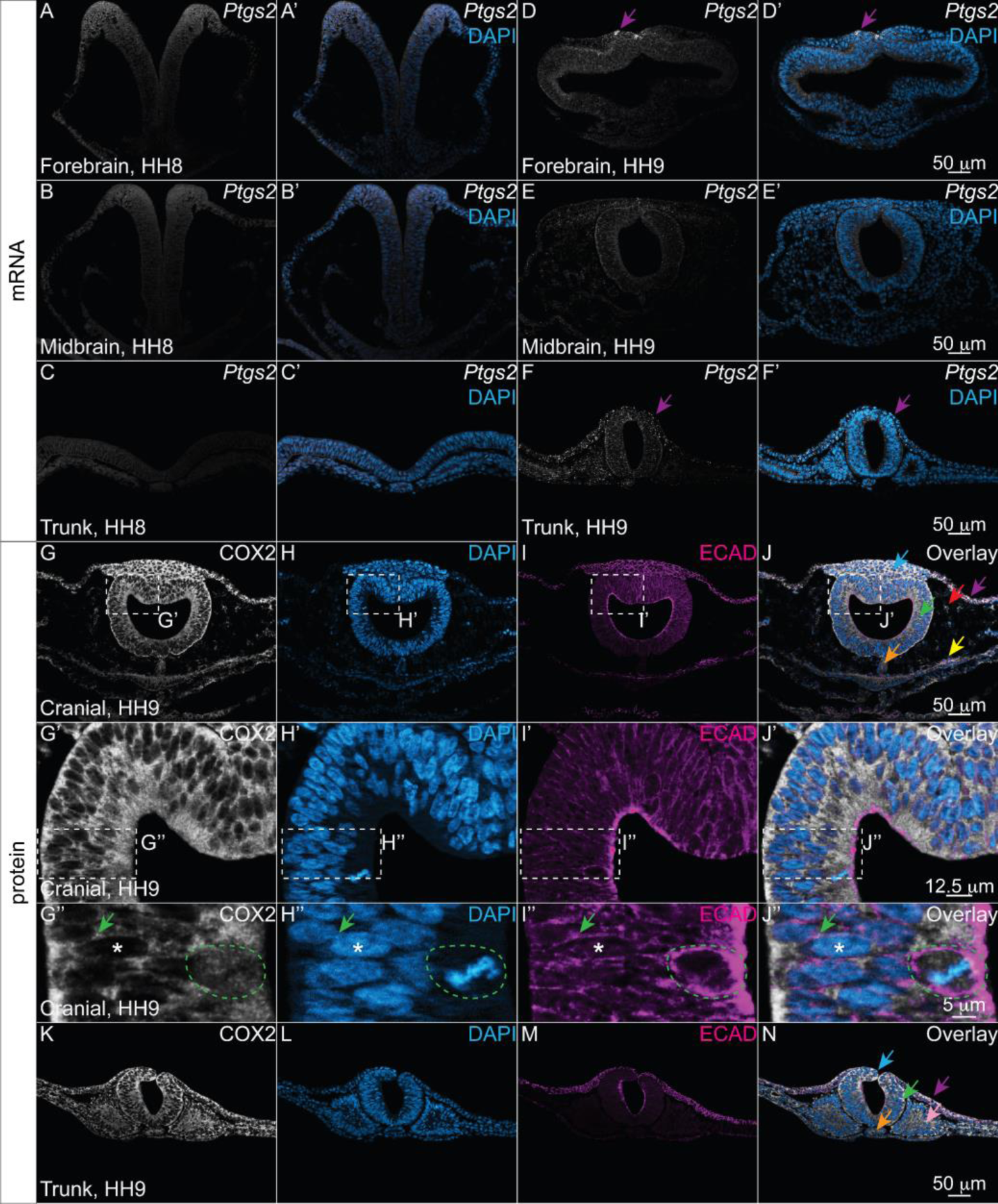
COX2 is broadly localized throughout the embryo during neurulation but its transcript is expressed at low levels. HCR for *Ptgs2* with DAPI stain at HH8 and HH9. (A-C’) Transverse cryosections of an HH8 chicken embryo and (D-F’) an HH9 chicken embryo from the forebrain, midbrain, and trunk axial levels showing COX2 transcript *Ptgs2* (white) and nuclear stain DAPI (blue). (D, D’, F, F’) Purple arrows indicate strong *Ptgs2* signal in the dorsal epidermis starting at HH9 in the forebrain and trunk regions. IHC for COX2 (white) with DAPI nuclear stain and ECAD (magenta). (G-N) Cranial and trunk transverse cryosections of an HH9 chicken embryo showing (G, G’, G’’, and K) the isoenzyme COX2 (white), (H, H’, H’’, and L) the nuclear stain DAPI (blue), and (I, I’, I’’, and M) ECAD (magenta). COX2 is expressed broadly throughout the embryo. (G”-J”) Within cells, COX2 signal colocalizes with ECAD (green arrow), is absent from DAPI+ nuclei (asterisk), and appears more diffuse in mitotic cells (green outline). Scale bar for each row in the last image of the row.

To characterize COX2 protein expression and localization, we performed IHC for COX2 with nuclear DAPI stain paired with membrane-localized epithelial cadherin (ECAD) in chicken embryos at stages HH7-10. We observed ubiquitous COX2 protein signal throughout the embryo (Figure 5G-5R). Within cells, COX2 localized to the cytoplasm and was absent from the nuclei marked by DAPI (Figure 5G’-5J’’, asterisk). In neuroepithelial cells, COX2 appears to overlap with ECAD in the lateral cell membranes (Figure 5G’’-5J’’, green arrow). In dividing cells, where the nuclear membrane has been dissolved, COX2 appears more diffuse within the cell (Figure 5G’’-5J’’, green outline). COX2 protein expression is similar in the trunk axial levels, with ubiquitous expression across cell types and protein localization to the cytosol (Figure 5K-5N).

### mPGES-2 protein and transcript expression and unique subcellular localization during neurulation

ScRNA-sequencing data showed that across all developmental stages included, the transcript encoding the terminal prostanoid synthase mPGES-2 was expressed widely across multiple tissues (Figure 1). mPGES-2 acts downstream of the COX isoenzymes to convert PGH_2_ into PGE_2_ (15). To date, no studies have characterized the spatial expression profiles of mPGES-2 or its corresponding gene *Ptges2* in the developing embryo. To characterize *Ptges2* expression in vertebrate embryos during neurulation, we used HCR in chicken embryos at stages HH7-10. We observed *Ptges2* expression in all tissues, but it appeared strongest in the neural tube, neural crest, and mesodermally derived tissues including the somites (Figure 6A-6F, green, blue, and pink arrows).

**Figure 6.**
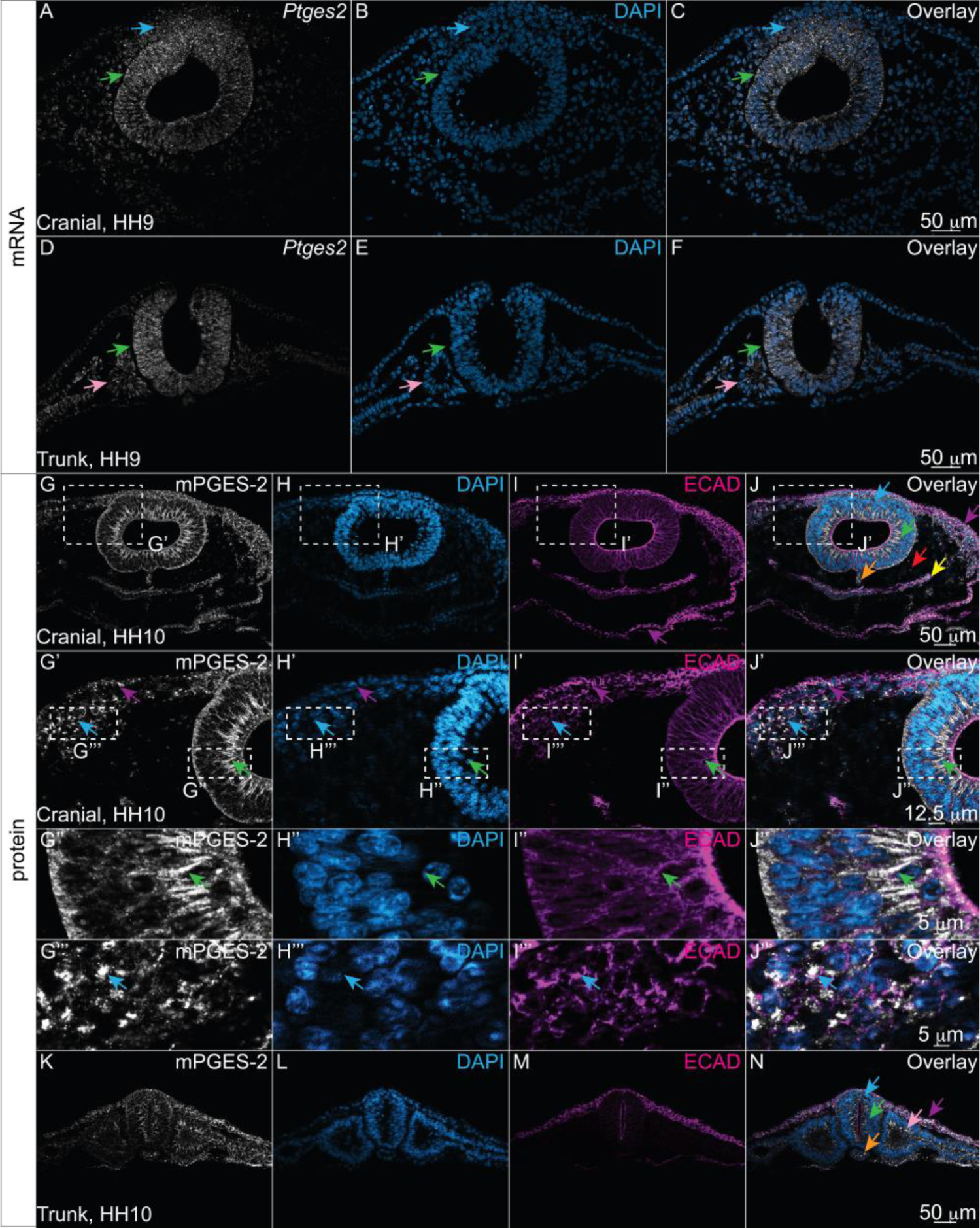
mPGES-2 protein and transcript are present in multiple cell types. HCR for *Ptges2* and DAPI nuclear stain. (A-F) Cranial and trunk transverse cryosections from an HH9 chicken embryo showing mPGES-2 transcript *Ptges2* (white) and the nuclear stain DAPI (blue). *Ptges2* signal appears most expressed in the neural tube, neural crest, and somite regions as indicated by green, blue, and pink arrows, respectively. (G-N) Cranial and trunk transverse sections of an HH10 chicken embryo with IHC showing the terminal prostanoid synthase mPGES-2 (white), nuclear stain DAPI (blue), and ECAD (magenta). mPGES-2 is broadly expressed across multiple tissues. (G’-J’) Zoom in of region outlined in (G-J) reveals that mPGES-2 co-localizes with ECAD in the membrane of neuroepithelial cells of the neural tube (green arrow), while in epidermal and neural crest cells, mPGES-2 is found in punctate compartments mutually exclusive from ECAD (purple and blue arrows, respectively). (G’’-J’’ and G’’’-J’’’) Zoom ins of G’-J’ in the neural tube and neural crest cells, respectively. Scale bar for each row in the last image of the row.

To characterize mPGES-2 protein during neurulation we performed IHC for mPGES-2 with DAPI nuclear stain and ECAD in chicken embryos at stage HH7-10. We observed mPGES-2 protein signal throughout the embryo across multiple cell and tissue types (Figure 6G-6N). In neuroepithelial cells, mPGES-2 co-localized with ECAD at the cell membrane (Figure 6G’-6J’’, green arrow). In contrast, mPGES-2 appeared localized to punctate condensates within epidermal and neural crest cells (Figure 6G’-6J’’’, purple and blue arrows, respectively). The protein was also detected in trunk axial level tissues and appeared to localize to the membrane with ECAD in epithelial epidermis, neural tube, and somites, but also showed punctate localization in a subset of cells (Figure 6K-6N). Of note, mPGES-2 is more distinctly visualized with 2% TCA fixation compared to 4% PFA fixation as observed with COX1 (Supp. Figure 3).

## Discussion

According to the classical understanding of cyclooxygenases, COX1 is the constitutive isoenzyme and COX2 is the inducible isoenzyme (22). This understanding appeared consistent at the transcript level in chicken embryos, where *Ptgs1* mRNA appeared ubiquitous while *Ptgs2* mRNA was spatiotemporally restricted (Figures 1, 3, 5). However, visualizing the corresponding protein localization demonstrated the importance of characterizing expression at both the gene and protein level.

Despite its broad gene expression, COX1 protein signal was only detectable in mitotic cells (Figure 3, 4). This stark contrast between mRNA and protein expression suggests that COX1 may be post-transcriptionally regulated to allow for dynamic changes in protein abundance depending on the cellular context. Past studies demonstrated that COX1 protein is degraded by the ubiquitin-proteasome system within ten minutes of intracellular calcium influx in human megakaryocytic MEG-01 cells (22). Cell cycle progression is also regulated by intracellular calcium, with intracellular calcium levels peaking at anaphase (32, 34), which correlates with the reduction of COX1 that we see in our mitotic cells. Our results suggest that in developing embryos, COX1 expression may be post-transcriptionally or post-translationally regulated depending on the cellular context and environment (Figure 4).

The specific localization of COX1 to mitotic cells suggests that COX1 may play a role in cell division. Past work identified that exposure to COX-inhibiting NSAIDs prevents cell division *in vitro* in multiple cancer cell lines (35, 36) and COX2 inhibition downregulates expression of genes associated with the spindle assembly checkpoint (37). Additionally, the signals and receptors downstream of COX1 and COX2 are linked with cell proliferation (38–40). Future research should investigate the role of COX isoenzymes, and particularly COX1, in cell division during embryogenesis.

In contrast to COX1, the *Ptgs2* gene was expressed sporadically at low levels and COX2 protein was detected ubiquitously throughout embryonic cell types during neurulation stages (Figure 1, 4). This break from previously described COX expression patterns may be attributed to the unique cellular context of embryonic development. During embryogenesis, drastic morphological changes occur on a cellular and tissue level. For example, neural crest cells undergo an epithelial to mesenchymal transition (EMT) whereby they delaminate from the neuroepithelial cells and migrate dorsolaterally out of the neural tube upon its fusion (41). While processes like EMT are normal and necessary in the embryo, they would represent a disease state in the adult. In fact, COX2 is upregulated in several cancers (42, 43) and drives cancer cell EMT and invasion (13, 44, 45).

Downstream of the COX isoenzymes, we characterized the expression of the terminal prostanoid synthase, mPGES-2. mPGES-2 functionally couples with both COX1 and COX2 to synthesize PGE_2_ (15). By synthesizing PGE_2_, mPGES-2 can have widespread effects as PGE_2_ is the most abundant PG in the adult and plays both homeostatic, pro-inflammatory, and anti-inflammatory roles depending on the context (24). Notably, PGE_2_ has implications in ovulation, cardiogenesis, and neural crest development based on prior research (1). In the neurulating chicken embryo, the *Ptges2* transcript was expressed broadly with increased expression in the neural tube and neural crest cells. The protein, mPGES-2, appeared to be expressed in all tissues that we visualized (Figure 6), which would theoretically allow it to work in conjunction with both ubiquitous COX2 and spatially restricted COX1 enzymes. Within cells, mPGES-2 varied in subcellular localization (Figure 6). It is reported to be synthesized as a Golgi membrane-associated protein and then localize to the cytosol in its mature form (15, 24). The varied subcellular localizations observed in different cell types could represent mPGES-2 in its various maturity states and their associated localizations (Figure 6). In addition, the dynamic subcellular localization of mPGES-2 was linked to different cell types. In neuroepithelial cells, the main mPGES-2 signal appeared in the cell membrane and co-localized with ECAD, but in the collectively migrating and mesenchymal migratory neural crest cells, mPGES-2 appeared to be localized to either subcellular compartments, vesicles, or condensates. Future work will focus on identifying how mPGES-2 subcellular localization may affect its function, specifically if membrane-localized mPGES-2 facilitates cell adhesion or morphological changes (e.g., neural tube closure) and if compartmentalization is necessary for neural crest migration.

In this study we have characterized the spatiotemporal expression of COX pathway enzymes COX1, COX2, and mPGES-2 in neurulating amniotic embryos. Notably, we found that the expression and localization of COX isoenzymes may not fit into previously defined expression patterns from disease cells and adult tissues in the embryo. The dynamic differences observed between transcript and protein signal highlights the need to characterize expression at both the gene and protein level to understand better when and where factors may be functioning in developmental processes. Moreover, we demonstrate the importance of validating antibodies in multiples fixatives (Supp. Figure 3). Future work will focus on defining the role of these COX pathway enzymes and the signals they produce during embryonic development.

## Methods

### Chicken Embryos

Fertilized chicken eggs were obtained from the Hopkins Avian Facility at the University of California, Davis and incubated at 37°C to the desired stages according to the criteria of Hamburger and Hamilton (HH). Use and experiments on embryos was approved by the University of California, Davis.

### Single cell analysis

COX pathway members in the *Gallus gallus* genome were identified using Ensembl Release 112. Expression profiles for each gene listed in Supp. Table 1 are shown in Supp. Figure 2. Filtered features matrices for embryos were obtained from peer-reviewed publicly available single cell datasets (NCBI GSE181577 (46) and GSE221188 (47)). Analyses were performed using Seurat V5 (48). Filtered feature matrices were independently log-normalized and scaled. Objects were integrated by sample using the harmony package (49) following which dimensionality reduction (determined using knee identification in elbow plot), k-means clustering (resolution determined using clustree package (50)) and neighbor identification was performed. The embeddings were utilized for Uniform Manifold Approximation and Projection (UMAP) plotting. Cell type annotations for clusters were performed using marker expression from the source manuscripts (46, 47). Kernel density gene expression plots were created using the Nebulosa package (51) and violin plots using the Seurat native VlnPlot function. Figures were organized in BioRender.

### Wholemount in situ hybridization chain reaction

Wholemount *in situ* hybridization chain reaction (HCR) was performed using the protocol suggested by Molecular Technologies with minor modifications. Chicken embryos were fixed in 4% paraformaldehyde in phosphate buffer (4% PFA) for 1 hr at room temperature (RT), washed in 1X PBS with 0.1% Triton (PBST), and were dehydrated in a series of 25%, 50%, 75%, and 100% methanol. Embryos were stored at -20°C prior to beginning HCR protocol. Embryos were rehydrated in a series of 25%, 50%, 75%, and 100% PBST when beginning the HCR procedure, but were not incubated with proteinase-K as suggested by the protocol. Embryos were incubated with 2.5-10uL of probes dissolved in hybridization buffer overnight (12-24 hr) at 37°C. After washes on the second day, embryos were incubated with 10uL each of hairpins H1 and H2 diluted in amplification buffer at RT overnight (12-24 hr). Embryos were subsequently incubated with 1:500 DAPI in PBST for 1 hr at RT, postfixed in 4% PFA for 1 hr at RT or 4°C overnight (12-24 hr), then washed in 1X PBS with 0.1% Tween-20 (PTween). After HCR, all embryos were imaged in both whole mount and transverse section (after embedding in gelatin and cryosectioning frozen samples) using a Zeiss Imager M2 with Apotome capability and Zen optical processing software.

### Immunohistochemistry

Immunohistochemistry (IHC) was performed as described previously and antibodies used in study are listed in Table 2 (33). Briefly, for IHC, chicken embryos were fixed on filter paper in 4% PFA for 15-20 min at RT or in 2% trichloroacetic acid (TCA) for 1 hr at RT. After fixation, embryos were washed in 1X TBS (500 mM Tris-HCl, pH 7.4, 1.5 M NaCl, and 10 mM CaCl_2_) containing 0.1% Triton X-100 (TBST+ Ca^2+^). For blocking, embryos were incubated in TBST+ Ca^2+^ with 10% donkey serum (blocking buffer) for 1 hr at RT or overnight (12-24 hr) at 4°C. Primary antibodies were diluted in blocking buffer at indicated dilutions and incubated with embryos for 48–96 hr at 4°C. After incubation with primary antibodies, whole embryos were washed in TBST+ Ca^2+^, then incubated with AlexaFluor secondary antibodies diluted in blocking buffer (1:500) overnight (12-24 hr) at 4°C. They were then washed in TBST+ Ca^2+^ and 4% PFA-fixed embryos were post-fixed in 4% PFA for 1 hr at RT. Antibodies used in the study (Table 2): Rabbit IgG α-COX1 (Cayman Chemical 160109), Rabbit IgG α-COX2 (Cayman Chemical 160126), Rabbit IgG α-mPGES-2 (Cayman Chemical 160145), Mouse IgG2a α-ECAD (BD Transduction Laboratories, 61081), and Mouse IgG1 α-Alpha Tubulin (Invitrogen, 322588). After IHC, all embryos were imaged in both whole mount and transverse section (after embedding in gelatin and cryosectioning frozen samples) using a Zeiss Imager M2 with Apotome capability and Zen optical processing software.

**Table 2.** Antibodies used in the study.

| **Antibody target** | **Dilution** | **Species Reactivity** | **Isotype/ Host** | **Immunogen** | **Source** | **Optimal Fixation (see Supp. Fig. 3)** |
| --- | --- | --- | --- | --- | --- | --- |
| Cyclooxygenase 1 (COX1) | 1:200 | Mouse, ovine | Rabbit IgG | Peptide from the internal region of mouse COX-1 | Cayman Chemical 160109, | 1h TCA |
| Cyclooxygenase 2 (COX2) | 1:200 | Human, macaque monkey, mouse, ovine, rat | Rabbit IgG | Synthetic peptide corresponding to the C-terminal region of mouse COX-2 | Cayman Chemical 160126, | 20m PFA |
| Microsomal Prostaglandin E Synthase-2 (mPGES-2) | 1:200 | Human, African green monkey, bovine, mouse, ovine, rat | Rabbit IgG | Synthetic peptide from the internal region of human mPGES-2 | Cayman Chemical 160145, | 1h TCA |
| ECAD | 1:500 | Human, Mouse, Rat, Dog | Mouse IgG2a | Amino acids 735-883 | BD Transduction | 20m PFA |
|  |  |  |  |  | Laborator<br>ies<br>61081, |  |
| Alpha tubulin | 1:200 | Human,<br>Mouse,<br>Rat, Sea<br>urchin | Mouse<br>IgG1 | Sarkosyl-<br>resistant filament<br>from sea urchin<br>sperm<br>axonemes | Invitroge<br>n,<br>322588 | 20m<br>PFA |

## 5. AUTHOR CONTRIBUTIONS

Conceptualization: CDR and TAL. Data Curation (experiments and imaging): TAL and RR. Formal Analysis: TAL and RR. Funding Acquisition: CDR and TAL. Methodology: CDR, TAL, and RR. Writing: CDR, TAL, and RR. Supervision: CDR.

## 6. ACKNOWLEDGEMENTS

The authors would like to acknowledge the following funding sources: NSF CAREER award 2143217 and NIH R03DE032047-01 to CDR. TAL was funded by the NIH 5T32GM007377-45 and an NSF GRFP. We would like to thank the members of the Rogers Lab at UC Davis and the UC Davis community for their discussions and input on this project.

## Supplementary material

**Supplemental Figure 1.**
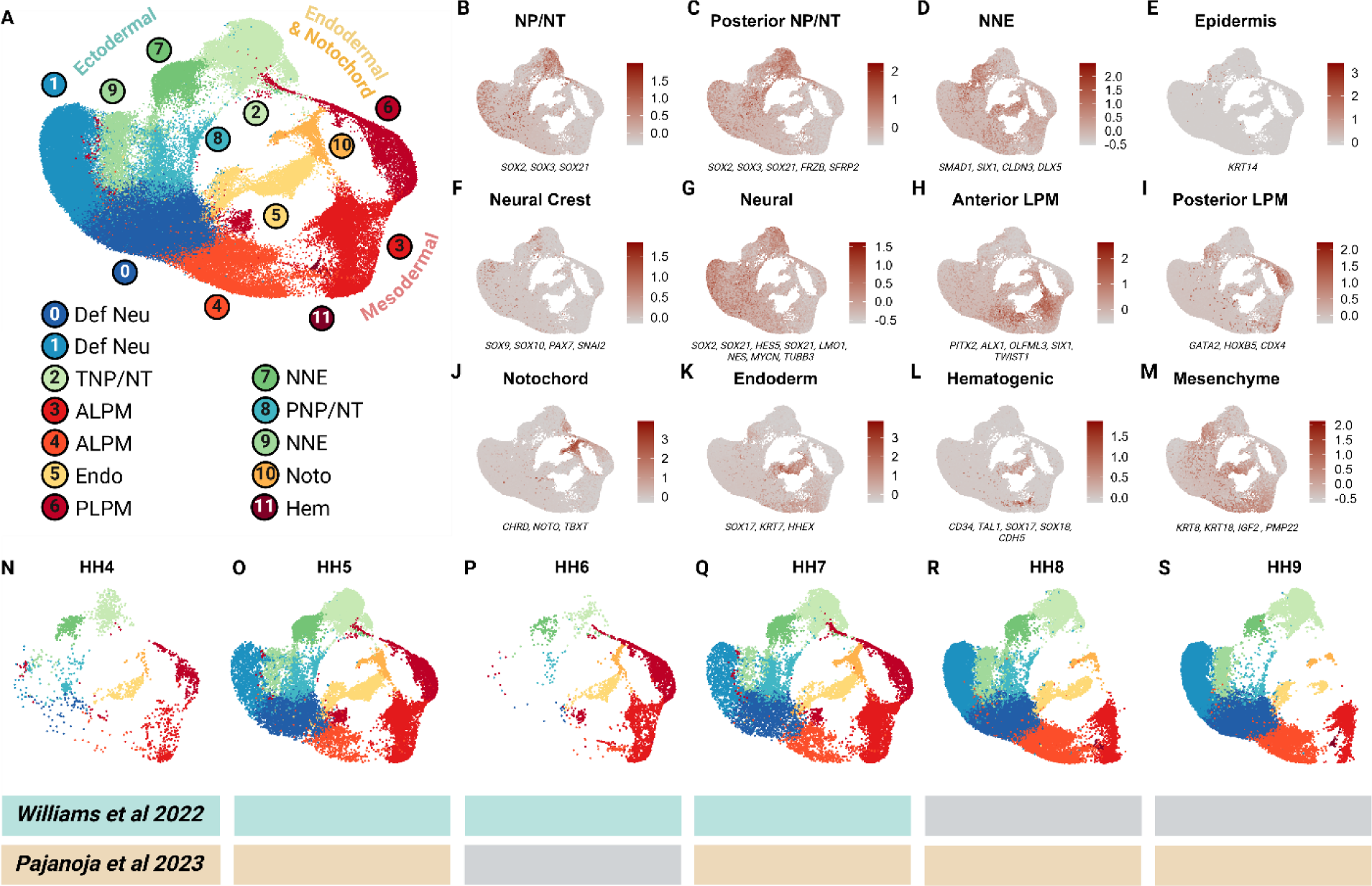
Combined clustering of scRNA-seq datasets to identify cell types. Unsupervised clustering of publicly available scRNA-seq data of chick embryos between HH4 and HH9 demonstrates the presence of 9 major cell types across 11 clusters. (A) UMAP demonstrating the unsupervised clustering results of chick whole embryos. (B-M) Feature plot expression of cell type marker gene modules. Genes included in each module are listed under each plot. (N-S) UMAP split by stages. Blue colored blocks indicate cells sourced from Williams *et al* 2022 and orange colored blocks indicate cells sourced from Pajanoja *et al* 2023. Grey blocks indicate lack of data from the source publication at the specific age. Definitive Neural lineage, Def Neu; Transitional neural plate/ neural tube, TNP/NT; Anterior lateral plate mesoderm, ALPM; Endoderm, Endo; Posterior lateral plate mesoderm, PLPM; Non-neural ectoderm, NNE; Posterior neural plate/ neural tube, PNP/NT; Notochord, Noto; Hematogenic cells, Hem.

**Supplemental Figure 2.**
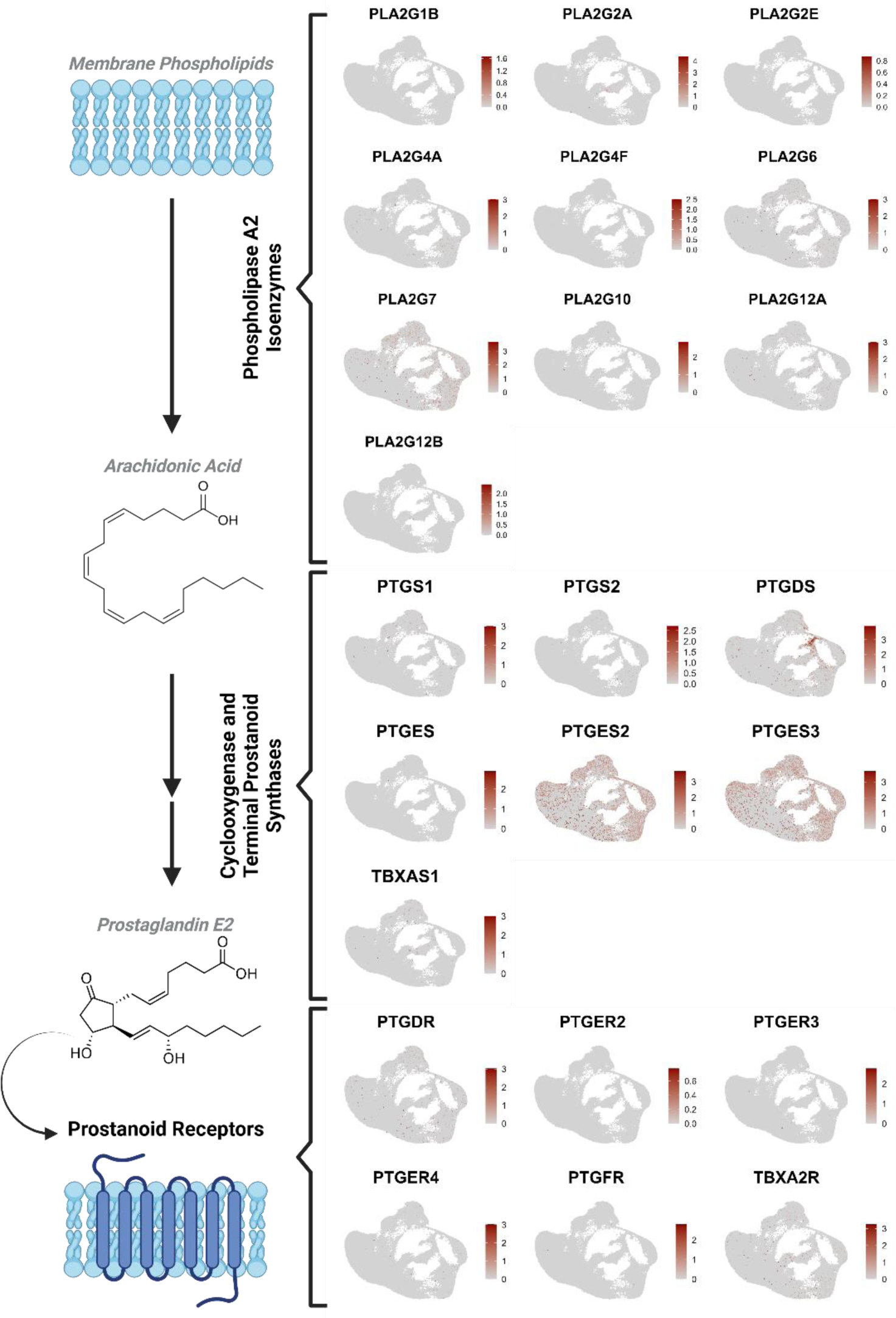
Expression of COX pathway members in publicly available scRNA-seq data of chick embryos between HH4 and HH9. (A-J) Feature plot demonstrating expression of phospholipase A2 isoenzymes. (K-Q) Feature plot demonstrating expression of cyclooxygenases and terminal prostanoid synthesis. (R-W) Feature plot demonstrating expression of prostanoid receptors.

**Supplemental Figure 3.**
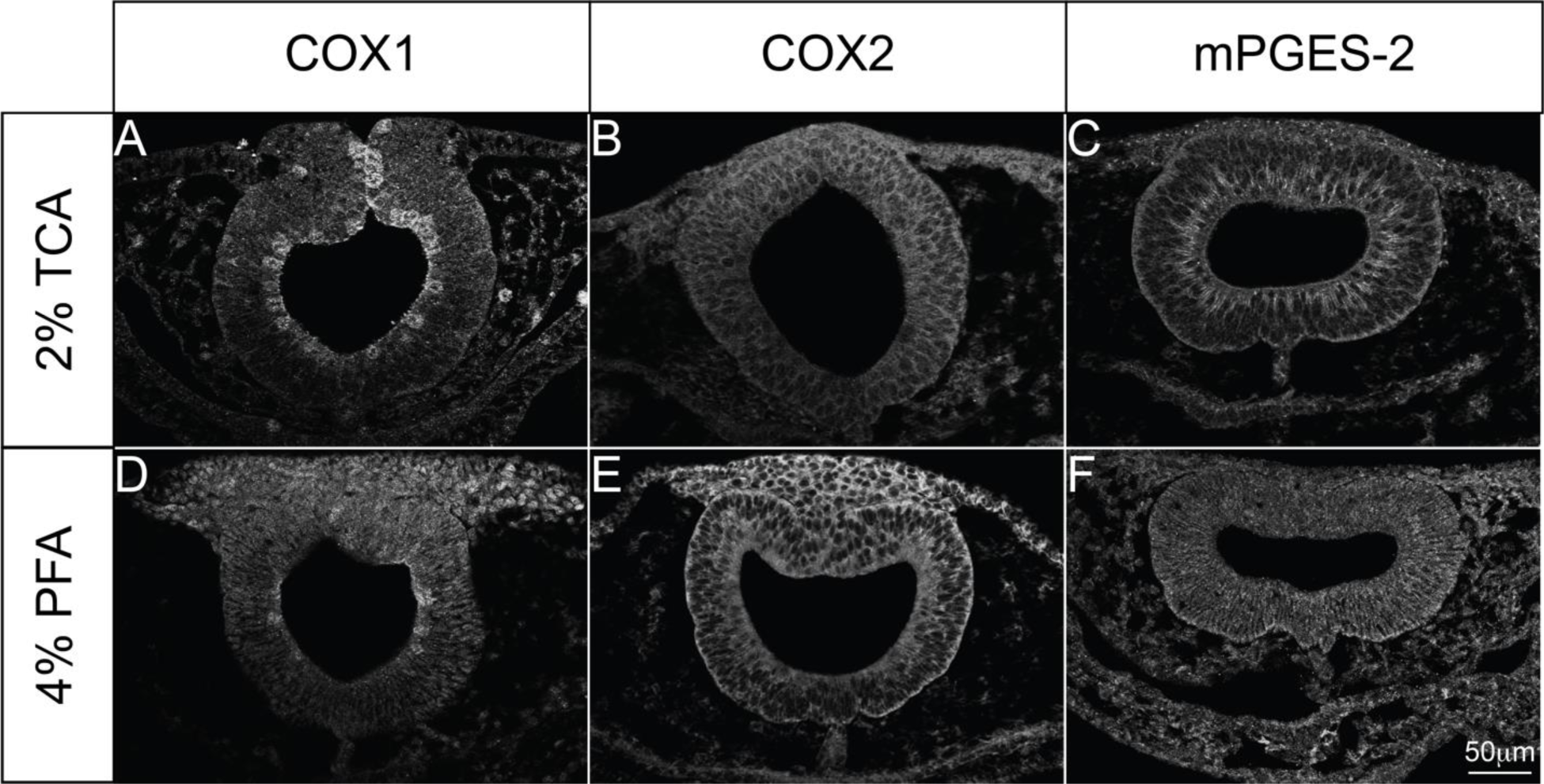
The type of fixation applied to the embryo affects IHC visualization. (A-C) Transverse sections of whole chicken embryos fixed with 2% TCA for 1hr. (D-F) Using the same antibodies as (A-C), transverse sections of whole chicken embryos fixed with 4% PFA for 15-20min with a 1hr 4% PFA postfix after antibody incubation. Protein visualization of (A) COX1 and (C) mPGES-2 appears more specific with 2% TCA fixation, while visualization of (B) COX2 appears clearer with 4% PFA fixation. Scale bar for all in (F).

**Supplemental Table 1.**
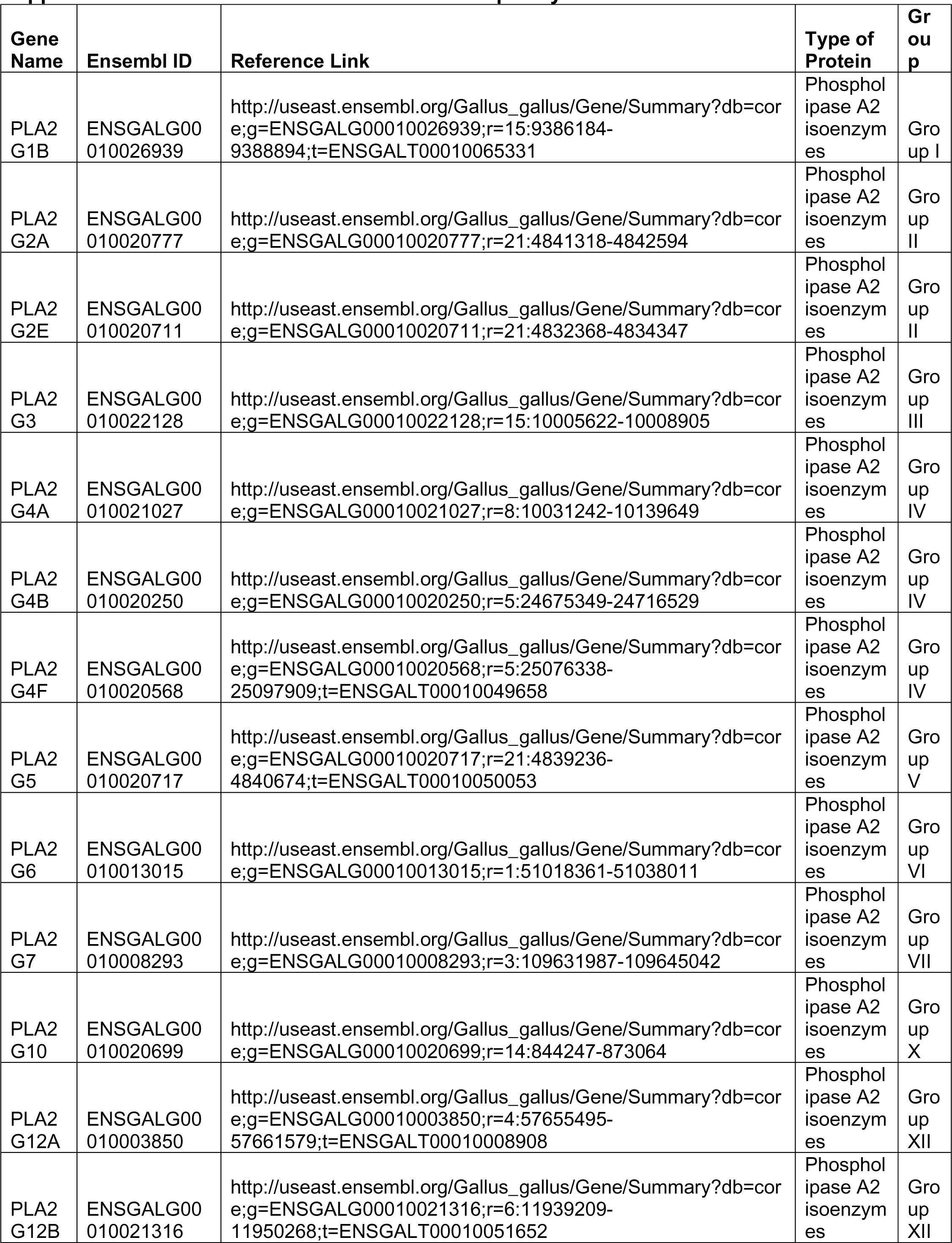
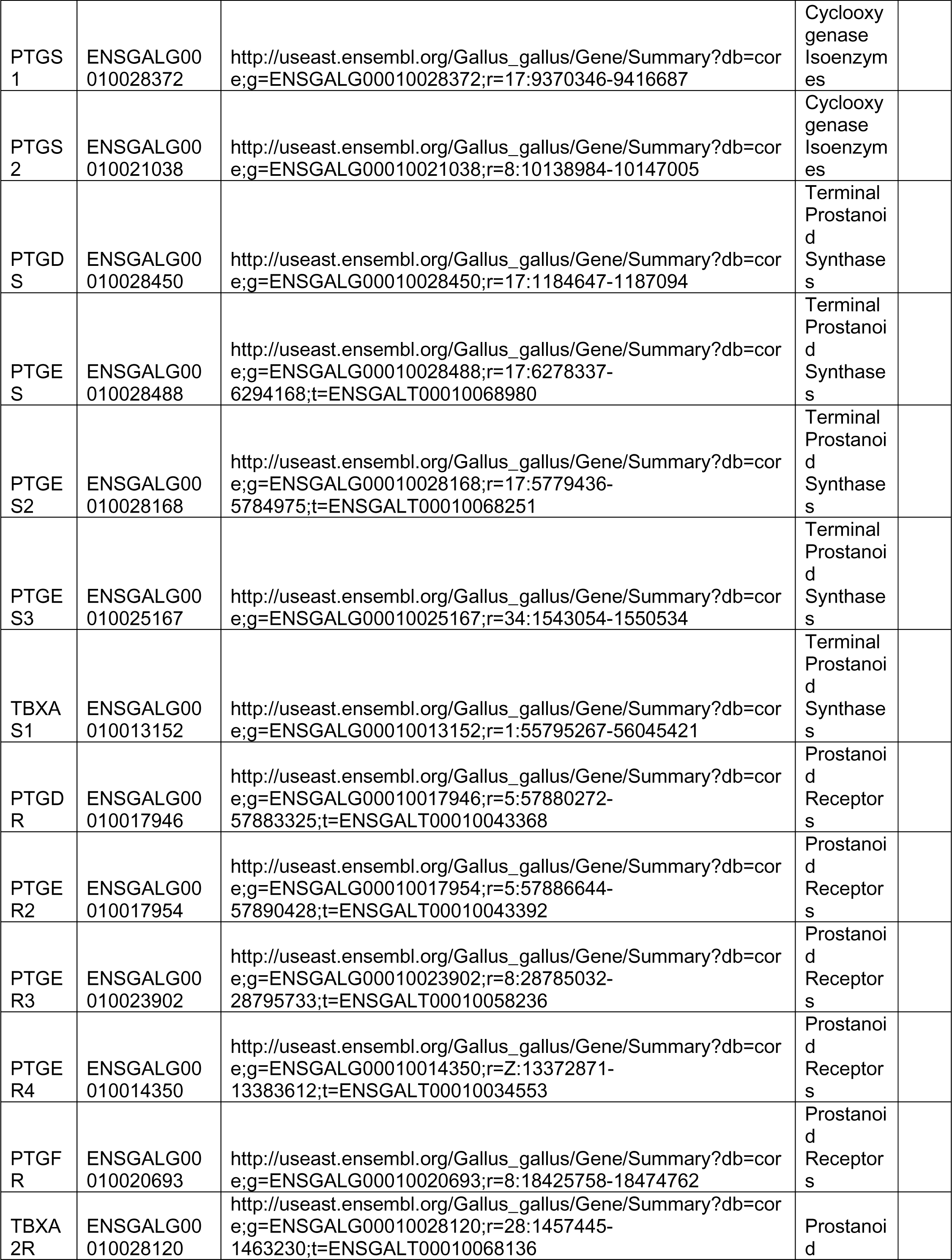
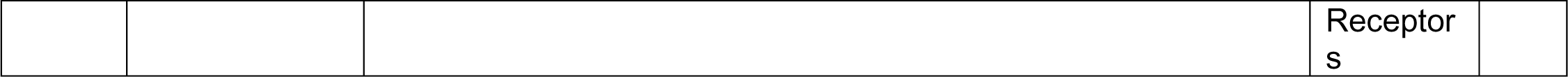
Genes used for scRNA-seq analysis.

